# Lethal streptococcal infection is associated with widened lymph node bottlenecks

**DOI:** 10.1101/2025.01.29.635438

**Authors:** Linya Xia, Karthik Hullahalli, Xingyu Tian, Chen Yuan, Fei Pan, Hongjie Fan, Matthew K. Waldor, Zhe Ma

## Abstract

*Streptococcus suis* serotype 2 (SS2) infection causes streptococcal toxic shock-like syndrome (STSLS), which can be lethal in both pigs and humans. In mouse models, relatively subtle differences in the inoculum size result in drastically different infection outcomes, from clearance to acute death. Here, we leveraged barcoded SS2 in a murine model to define the bacterial population dynamics associated with death. Barcode sequencing analysis demonstrated that a 10-fold increase in inoculum size led to a 100-1,000-fold increase in the number of bacteria that initiated infection in certain lymph nodes, ultimately driving lethality. This lethal dose threshold was in part driven by suilysin (SLY), the major SS2 virulence factor. At higher doses, the clearance and replication kinetics of bacteria lacking SLY mimicked WT SS2 at sub-lethal doses. Our findings suggest that acute death in SS2 infection is caused by a wholesale failure of the immune system, enabling a large number of bacteria to initiate infection. These results demonstrate how subtle changes in inoculum size can have dramatic consequences for infection outcome and how lethal infection is driven by widened infection bottlenecks.

**Author Summary:** Lethal infections are often associated with a specific dose threshold, where increasing inoculum size is associated with an increased likelihood of death. However, a quantitative explanation how this numeric threshold is established is lacking. For example, lethal dose thresholds may reflect a dose where at least one microbe can pass through host-imposed barriers to infection. Alternatively, lethal dose thresholds may reflect a failure of the host immune system, leading to numerous microbes initiating infection. Here, using barcoded *Streptococcus suis* serotype 2 (SS2), we compare the within-host population dynamics associated between lethal and non-lethal infections. We find that lethal doses are associated with dramatic widening of infection bottlenecks, leading to a large number of bacteria that initiate infection. Lethal infection is also driven by suilysin, the major SS2 virulence factor. These observations demonstrate how lethal doses are quantitatively established by the relationship between host bottlenecks and bacterial virulence factors.

## Introduction

*Streptococcus suis* serotype 2 (SS2) is an opportunistic zoonotic pathogen. SS2 infections are widespread in humans and pigs, with notably high morbidity and mortality rates reported in Asian and European countries [1-4]. Hosts succumb to infection due to streptococcal toxic shock-like syndrome (STSLS), which is driven by the expression of suilysin, the major SS2 virulence factor [5, 6]. STSLS is characterized by cytokine storm along with high bacterial load, organ dysfunction, and acute host mortality [7, 8].

In mice, a 10^7^ CFU infectious dose is cleared with no clinical signs while a 10^8^ CFU dose is lethal in 100% of animals within 8 hours. Higher doses have long been appreciated to lead to worse infection outcomes, yet a quantitative explanation for why some doses are lethal while others are cleared, and which host and bacterial factors contribute to the lethal dose range, is lacking. Given the dramatic difference in infection outcome between 10^7^ and 10^8^ challenge doses, this model therefore represents a powerful system to study how quantitative differences in pathogen populations confer lethality. Here, we leveraged barcoded bacteria to monitor the within-host population dynamics during SS2 infection in mice and investigated how relatively small changes in the infectious dose can lead to rapid clearance or death. We demonstrate high-dose SS2 infection results in marked pathology and dramatically widened infection bottlenecks in lymph nodes, potentially driving lethality.

## Results

### A 10-fold change in dose distinguishes rapid pathogen clearance and acute death

We leveraged in IP inoculation model to investigate how dose controls lethal infection outcomes in murine SS2 infection [9]. We found that SS2 infection results in drastically different survival phenotypes with challenge doses of 10^7^ and 10^8^ CFU (Fig. 1a). At 10^7^ CFU, all animals survived, while infection with 10^8^ CFU led to acute death by 8 hours post inoculation (Fig. 1a). There was also a marked difference in the CFU burdens between mice challenged with 10^7^ and 10^8^ CFU. We observed that mice challenged with a lethal dose (10^8^ CFU) had a greater CFU burden in most organs and lymph nodes at each time point within the first 7 hr post inoculation compared to those challenged with a non-lethal dose (10^7^ CFU) (Fig. S1b, s1c). In non-lethal infection, CFU burdens were reduced at 24 hours post infection and largely cleared by one week (Fig. S1b, s1c). In contrast, in all tissues from the lethal dose, bacterial burdens increased up to 7 hours, after which all mice died.

**Figure 1:**
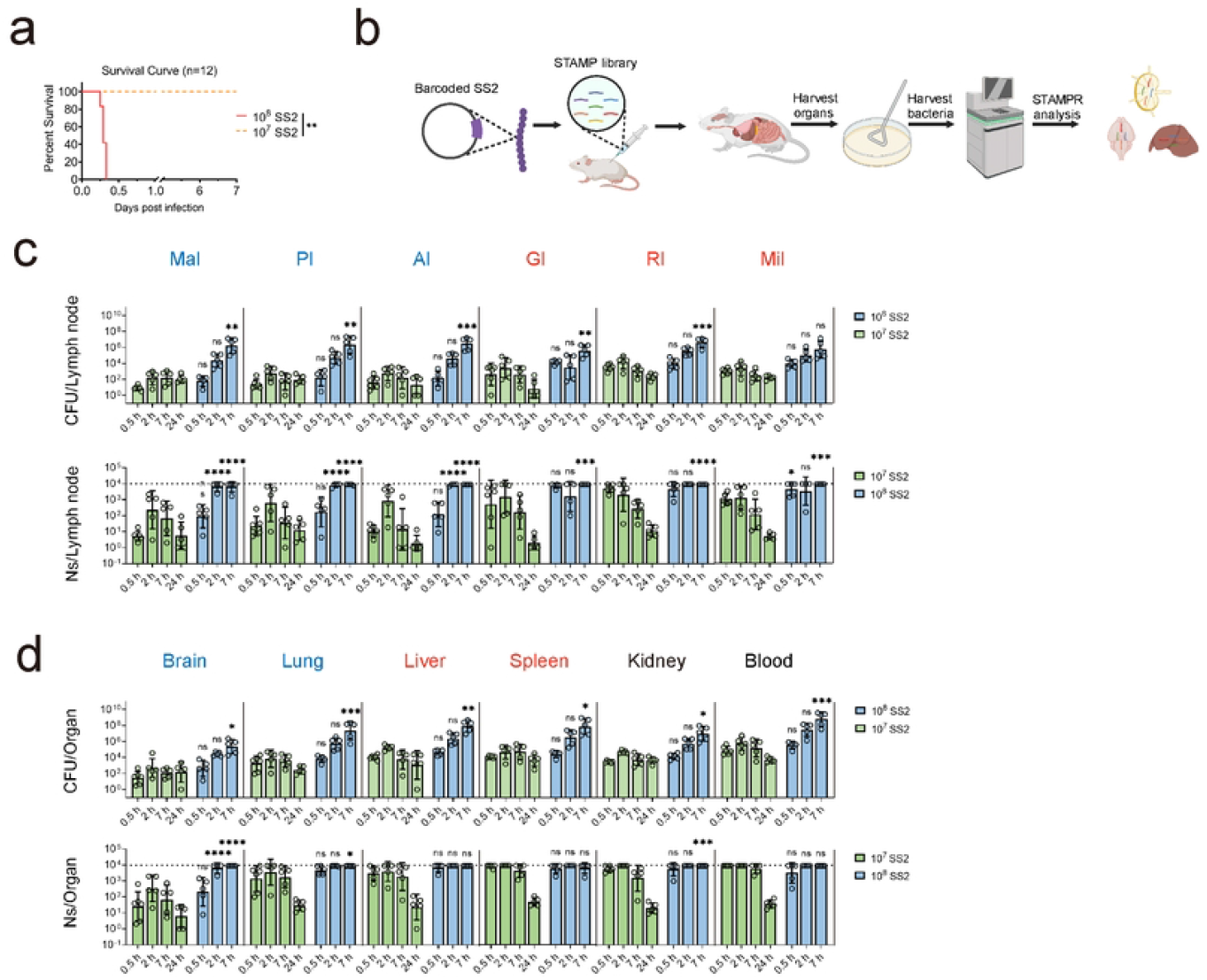
Lethal infection is associated with wider bottlenecks. (a) Survival curves of mice challenged with different doses (1×10^7^ CFU, 1×10^8^ CFU) of SS2. All mice challenged with 10^7^ CFU survived for at least 7 days, while those challenged with 10^8^ CFU experienced acute death within 8 hpi. (b) Schematic of the experimental protocol using the barcoded SS2 library. Each colored bacterium possesses a unique barcode. Mice were intraperitoneally injected with the barcoded library, and organs were harvested following cardiac perfusion, homogenized, and plated for enumeration of CFU. Bacteria were harvested, and a STAMPR analysis was performed. (c and d) The bacterial burden and Ns (FP size) in extraperitoneal (blue) lymph nodes (c) and organs (d), and intraperitoneal (red) lymph nodes (c) and organs (d) of mice. Mice were challenged with 10^7^ CFU of SS2_-STAMP_ library (Green squares) at 0.5 hpi (n=6), 2 hpi (n=5), 7 hpi (n=5), 24 hpi (n=5) and 10^8^ CFU of SS2_-STAMP_ library (Blue squares) at 0.5 hpi (n=5), 2 hpi (n=5), 7 hpi (n=5). Dotted line at 10^4^ indicate the resolution limit for Ns calculation, which represents the maximum possible FP size that can be quantified with this library. Bars indicate geometric means with geometric SD. Statistical significance was calculated using two-way ANOVA (c and d) followed by t-tests with Bonferroni correction between 10^7^ and 10^8^ CFU groups at the same time points: \*\*\*\**p* < 0.0001; ***p < 0.001; **p < 0.01; *p < 0.1, ns indicates no significant difference. Survival curves were analyzed using the Log-rank (Mantel-Cox) test for (a): \*\**p* < 0.01.

Both gross pathology and histopathological analyses revealed that the principal difference between lethal and non-lethal doses were apparent at 7 hpi in the mandibular (Mal) and parotid (Pl) lymph nodes. We observed notable erythema and swelling only at the lethal dose (Fig. S2a, s2c), along with severe congestion and the apparent infiltration of mononuclear phagocytes and lymphocytes in the cortex of the lymph node (Fig. S2b). Together, these results demonstrate that the increase in dose from 10^7^ to 10^8^, which distinguishes clearance and acute death, is associated with higher CFU burdens in all tissues and major pathology in two lymph nodes.

### Lethal infection is associated with wider bottlenecks

We reasoned that two distinct processes may drive the higher CFU burden observed in all tissues with the lethal 10^8^ dose. The higher bacterial loads at the lethal dose may be due to an increase in pathogen replication. Alternatively, the higher bacterial loads may be driven by a defect in ability of the host to eliminate inoculated microorganisms, or a combination of both processes. The relative contribution of clearance and replication can be assessed by quantifying the founding population (FP) size, which is defined as the number of cells from the inoculum that give rise to the bacterial population at the site of infection. Higher FP sizes relative to the dose reflect wider infection bottlenecks.

To quantify the infection bottlenecks associated with lethal and non-lethal doses of SS2, we generated a barcoded SS2 library with 919 unique barcodes and the STAMPR pipeline. This library can calculate FP sizes up to ∼10^4^ founders using the Ns metric. (Fig. S3e). At a lethal dose of 10^8^, Ns values increased from 0.5 to 2 hpi in the lymph nodes, indicating that bacterial clones continue to seed these tissues throughout the first few hours of infection. By 2 hpi, the Ns in most organs and lymph nodes had reached the resolution limit of the library (∼10^4^) and remained at least at that level until death at 7 hpi (Fig. 1c, 1d). In contrast, in mice challenged with a non-lethal dose of 10^7^, Ns in the lymph nodes peaked at 2 hpi, and then began to decline throughout the first 24 hours. Notably, only 1-10 founders were detected in lymph nodes at 24 hpi, a 6-log reduction relative to the inoculum size (Fig. 1c, 1d). At 7 hpi, which represents the latest time point prior to death in lethal infection, we observed that the 10-fold increase in dose was associated with 100-1000-fold increase in Ns across several lymph nodes. This marked widening of the infection bottleneck was particularly notable in the Mal and Pl, which displayed the most severe pathology.

By 24 hpi, Ns values were very similar to CFU values in the non-lethal model, indicating that each founder had undergone minimal net replication, reflecting our findings that SS2 is largely cleared by 1-week post infection. Together, these observations indicate that the 10^8^ dose is associated with higher FP in most organs. At a 10^7^ dose, fewer bacteria survive host defense mechanisms and consequently undergo little expansion. The magnitude of increase in the FP is notable, where a 10-fold increase in the dose led to a 100 to 1,000-fold increase in the FP sizes in the lymph nodes. Thus, lethal infection is associated with a dramatic widening of the infection bottleneck. These observations suggest that death is associated with a large number of SS2 clones breaching host barriers. In non-lethal infection, the number of SS2 cells in the host is apparently insufficient to damage host barriers, ultimately enabling successful clearance. The observation that the widening of the bottleneck is tissue-dependent suggests that certain lymph nodes in particular may play a crucial role in preventing death during SS2 infection.

### Suilysin controls the lethal dose threshold

Suilysin (SLY) is a major virulence factor in SS2 that contributes to the development of STSLS [6, 9]. In contrast to WT SS2, 10^8^ CFU of ΔSLY led to no mortality (Fig. S4d). We hypothesized that, like non-lethal infection in WT SS2, ΔSLY also experience tighter infection bottlenecks. Therefore, we constructed barcoded library in the *sly* gene deletion mutant (ΔSLY_-STAMP_) containing approximately 952 barcodes, which had a resolution limit up to ∼10^4^ founders. At 0.5 hpi, ΔSLY achieved CFU burdens comparable to WT in lymph nodes and organs. However, there was no increase in ΔSLY burden in most organs, in marked contrast to WT SS2, where CFUs increased until death (Fig. 2a, 2b). Ns values were also significantly lower at 7 and 24 hpi in ΔSLY (Fig. 2a, 2b), indicating that the clones that initially initiated infection were incapable of sustained colonization, similar to observations in WT SS2 at the sub-lethal dose. In contrast to WT SS2, in the Mal and Pl, Ns values did not increase over the first two hours in ΔSLY and ultimately declined by 24 hpi. These observations demonstrate that SLY enables SS2 to evade host bottlenecks to cause death.

**Figure 2:**
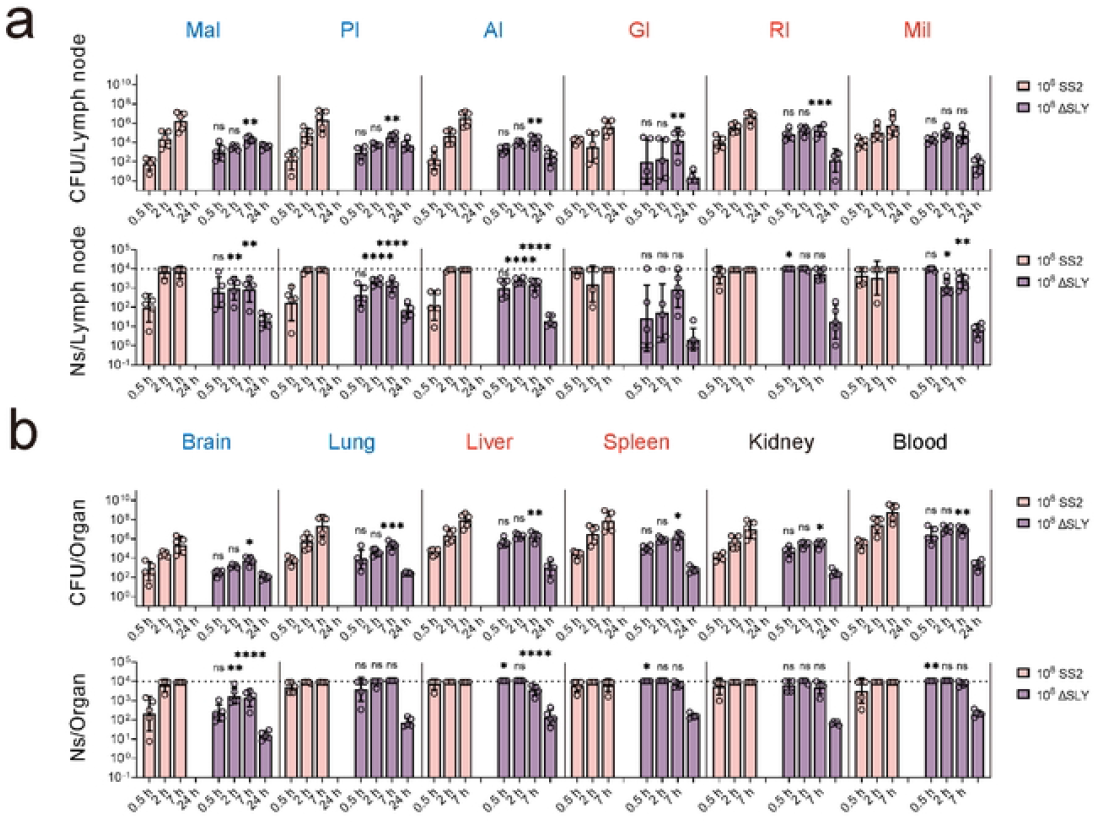
Suilysin controls the lethal dose. Bacterial burden and FP sizes (Ns) in extraperitoneal (blue) lymph nodes (a) and organs (b), and intraperitoneal (red) lymph nodes (a) and organs (b) of mice were measured after challenging with 10^8^ CFU of SS2_-STAMP_ library at 0.5 hpi (n=5), 2 hpi (n=5), 7 hpi (n=5), or 10^8^ CFU of ΔSLY_-STAMP_ library at 0.5 hpi (n=5), 2 hpi (n=5), 7 hpi (n=5) and 24 hpi (n=5). Dotted line at 10^4^ represents the resolution limit of the library. Bars indicate geometric means with geometric SD. Statistical significance was calculated using two-way ANOVA (c and d) followed by t-tests with Bonferroni correction between 10^7^ and 10^8^ CFU groups at the same time points: \*\*\*\**p* < 0.0001; ***p < 0.001; **p < 0.01; *p < 0.1, ns indicates no significant difference.

Elevated CRP levels were observed only in mice infected with the WT strain at 10^8^ CFU, while ΔSLY and WT at 10^7^ CFU did not trigger CRP expression (Fig. S5). These results highlight the role of SLY in evading host defenses through excessive inflammatory responses [10], ultimately leading to lethal outcomes.

## Discussion

Our study utilizes barcoded SS2 to reveal that lethal infection involves a significant widening of the infection bottleneck in specific lymph nodes. Small changes in inoculum size or vital virulent factors function can drastically alter infection outcomes, emphasizing the role of bottlenecks in preventing lethal infections.

We observe that SLY is required for lethal infection and in part controls the lethal dose threshold. At 10^8^ CFU, WT SS2 is lethal and ΔSLY is not. Since it is likely that bacterial surface molecules sensed by the immune system are similarly abundant between WT and ΔSLY, the difference in lethality likely indicates that the ability of SS2 to drive excessive inflammatory responses at 10^8^ CFU is not driven by innate sensing of bacterial surface molecules. Rather, the difference in lethality is likely attributed to a specific function of suilysin, such as direct killing or inactivation of immune cells. These observations are significant since they suggest that the host immune factors that control the lethal dose threshold in SS2 infection are unlikely to be processes that sense extracellular bacteria (e.g., TLRs), and instead are potentially other factors that confer sensitivity to suilysin-dependent killing. Future work can perturb these host factors to decipher the molecular mechanisms that underlie lethal dose thresholds.

Unexpectedly, we observed significant pathology in the Mal and Pl but not at other sites. These sites also exhibited the greatest widening of infection bottlenecks between lethal (10^8^ CFU WT) and non-lethal (10^7^ CFU WT or 10^8^ CFU ΔSLY) models. We speculate these observations reflect the importance of these specific lymph nodes in controlling SS2 infection, perhaps by limiting bacterial dissemination into the brain. The importance of lymph nodes in controlling streptococcal infection has also been observed in a model of *S. pyogenes* systemic infection [11]. More broadly, the application of barcoded bacteria to study how lymph nodes controlling bacterial dissemination will help further elucidate host mechanisms of infection control.

We previously developed a nomenclature in a model of *E. coli* systemic infection at a non-lethal dose range [12]. Our findings from SS2 lethal infection suggests that “negative scaling” (the widening of bottlenecks at higher doses) occurs around the lethal dose threshold. While the molecular mechanisms behind bottlenecks vary across infection contexts, it is plausible that these mechanisms share similar quantitative traits. If generalizable, lethal infectious doses might correspond to the range where bottlenecks fail to scale effectively.

## Material and Methods

### Bacterial Strains and Growth

The SS2 strain ZY05719 was cultured in Todd-Hewitt Broth (THB, Becton, USA). E. coli was cultured in Lysogeny broth (LB, OXOID, UK) at 37 °C. Additional compounds and antibiotics were used at the following concentrations: Spectinomycin (Spc, 50 μg/mL), Kanamycin (Kan, 50 μg/ml). Further details regarding construction of bacterial strains and animal experiments are provided in *Supporting_Protocol*.

### STAMP procedure

Briefly, mice were injected with barcoded libraries, and organs were harvested and plated for enumeration of CFU. Bacteria were harvested and sequenced with a NovaSeq (Illumina), and STAMPR analysis was performed as previously described [13, 14]. Further details regarding construction and analysis of barcoding experiments are provided in *Supporting_Protocol*.

## Acknowledgments

This research was supported by the National Key Research and Development Program of China (2021YFD1800800); the National Natural Science Foundation of China (32273009); the Fundamental Research Funds for the Central Universities (KJJQ2024005); the National Key Research and Development Program of China (2021YFD1800404); Jiangsu Agriculture Science and Technology Innovation Fund (CX(23)1041); the Priority Academic Program Development of Jiangsu Higher Education Institutions; and the “Young Scholars” cultivation program of College of Veterinary Medicine in Nanjing Agricultural University; Postgraduate Research and Practice Innovation Program of Jiangsu Province (KYCX24_0996). The authors thank Zhong Tang (College of resources and environmental sciences, Nanjing Agricultural University) for assistance in barcode sequencing.

## Supplementary Figure Captions

**Figure S1:**
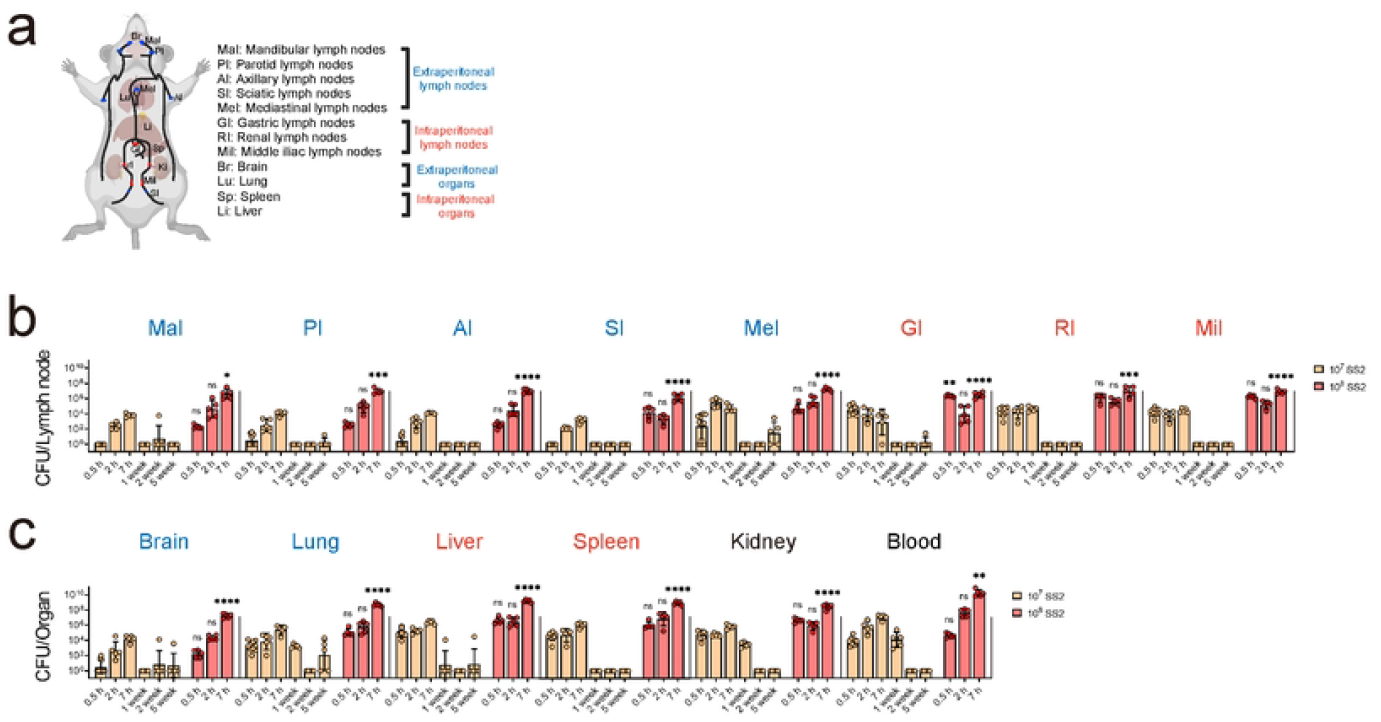
The CFU burdens of lethal and non-lethal infections of SS2. (a) After intraperitoneal infection with SS2 in BALB/c mice, organs were collected following cardiac perfusion, including the extraperitoneal lymph nodes (Mal, Pl), Axillary lymph nodes, Sciatic lymph nodes, Mediastinal lymph nodes, marked with blue circle and blue text), the intraperitoneal lymph nodes (Gastric lymph nodes, Renal lymph nodes, Middle iliac lymph nodes, marked with red circle and red text), the extraperitoneal organs (Brain, Lung) and intraperitoneal organs (Spleen, Liver). (b and c) Bacterial burden of the lymph nodes (b) and organs (c) challenged with 1×10^7^ CFU (Left, Orange) at 0.5 hpi (n=10), 2 hpi (n=5), 7 hpi (n=5), 1 week post infection (n=5), 2 weeks post infection (n=5), 5 weeks post infection (n=5) and 1×10^8^ CFU (Right, Red) of SS2 at 0.5 hpi (n=5), 2 hpi (n=5), 7 hpi (n=5). Bars indicate geometric means with geometric SD. Extraperitoneal lymph nodes and organs were marked with blue text, and intraperitoneal lymph nodes and organs were marked with red text. Statistical significance was calculated using two-way ANOVA for (b and c) for 10^7^ SS2 group and 10^8^ SS2 group at 0.5 hpi, 2 hpi, 7 hpi. Two-way ANOVA analysis is generally followed up by Bonferroni test to identify statistic: \*\*\*\**p* < 0.0001; \*\*\**p* < 0.001; \*\**p* < 0.01; \**p* < 0.1, ns indicates no significant difference.

**Figure S2:**
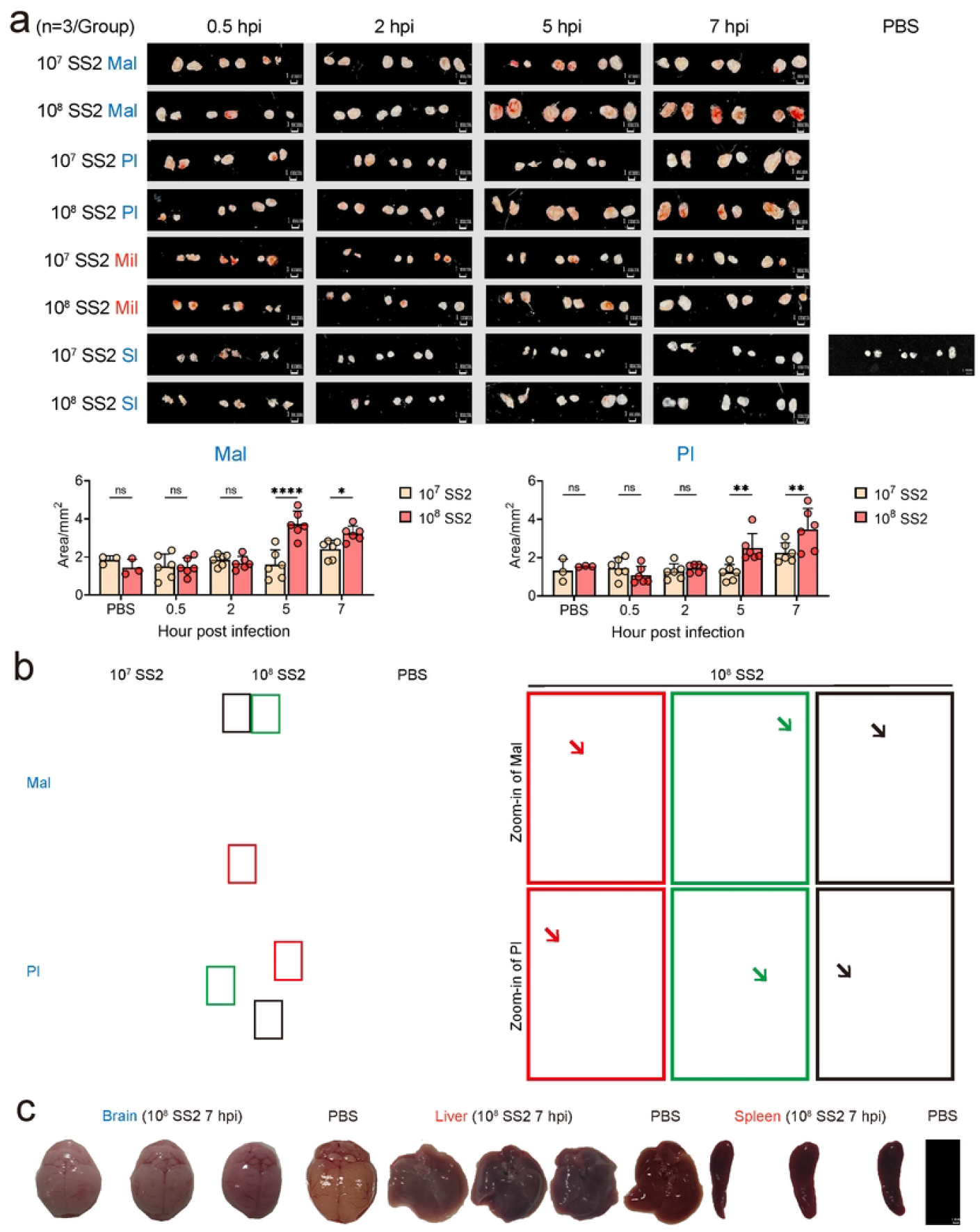
SS2 infection leads to severe pathology in the mandibular (Mal) and parotid (Pl) lymph nodes. (a) Mal, Pl, Mil, Sl were harvested at different time points (0.5 hpi, 2 hpi, 5 hpi, 7 hpi, n=3). After intraperitoneal infection with a lethal dose of SS2, visible lesions and swelling were observed in Mal and Pl at 5 hpi and 7 hpi. Images were analyzed using ImageJ. (b) Hematoxylin and eosin (H&E) staining of pathological sections of Mal and Pl from mice infected with SS2 or PBS control at 7 hpi. Zoom-ins are indicated by red (severe congestion), green (infiltration of lymphocytes in the cortex of the lymph node), and black (infiltration of mononuclear phagocytes) squares. (c) The brain, liver, and spleen were collected at 7 hpi from mice infected with a lethal dose of SS2, and no visible lesions were observed (n=3). Statistical significance was calculated using two-way ANOVA for (a): \*\*\*\**p* < 0.0001; \*\**p* < 0.01; \**p* < 0.1, ns indicates no significant difference.

**Figure S3:**
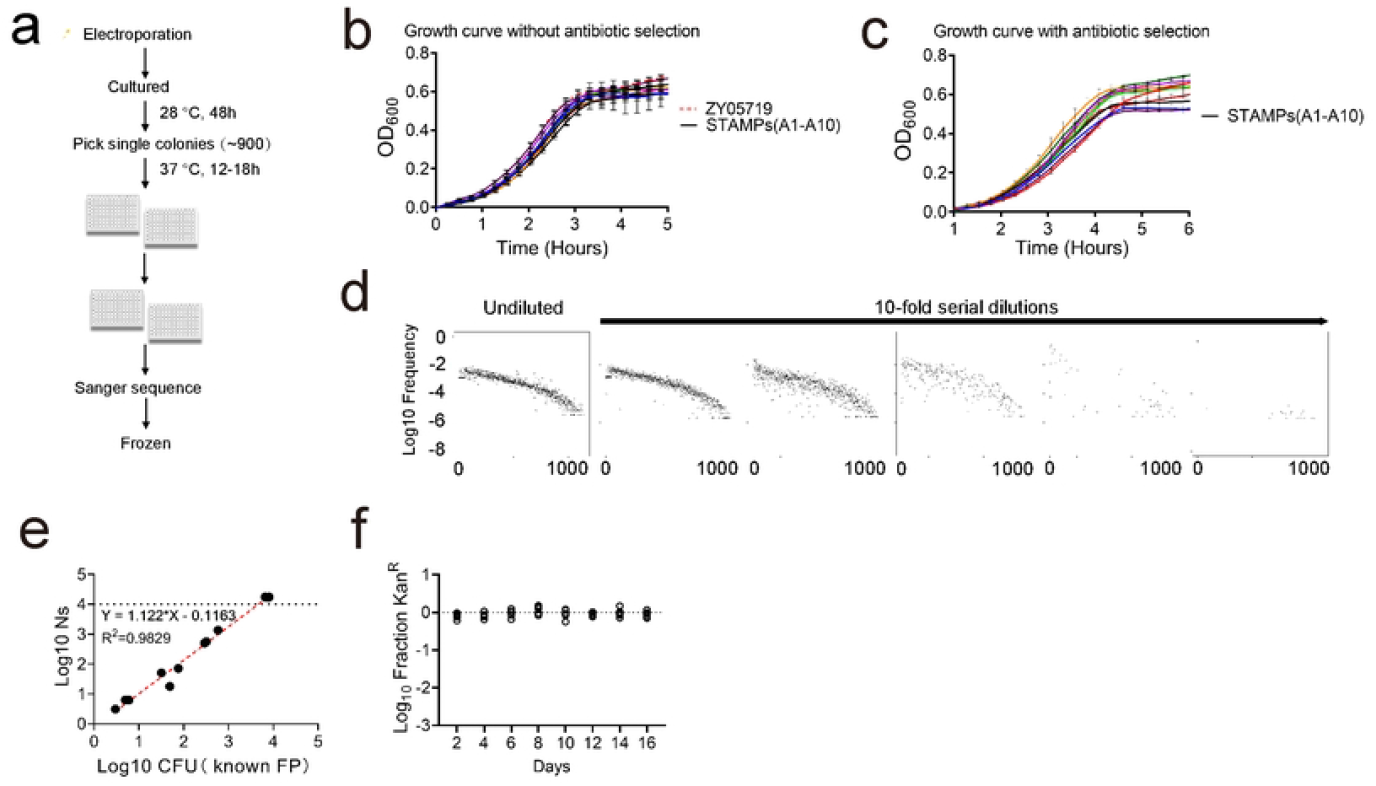
SS2 STAMP library construction. (a) Experimental schematic for the generation of the SS2 barcoded library. Individually barcoded bacteria were inoculated in single well of 96-well plates. ∼900 individual colonies were pooled and frozen. (b, c) Growth curves of 10 individual barcoded clones and WT in THB medium without Kan (b) or with Kan (c). The barcode insertion did not cause a growth defect. (d, e) Sequencing of the barcodes at various known bottleneck sizes from serial dilution (in vitro bottlenecks) demonstrated that Ns accurately reflects FP sizes up to 10^4^. (f) The barcoded library was serially passaged in THB medium without Kan. Every two days, the fraction of cells containing barcodes were quantified as the fraction of kanamycin-resistant CFUs. All colonies retained the antibiotic resistance marker linked to barcode, indicating that the barcode is stable for at least 16 days without kanamycin.

**Figure S4:**
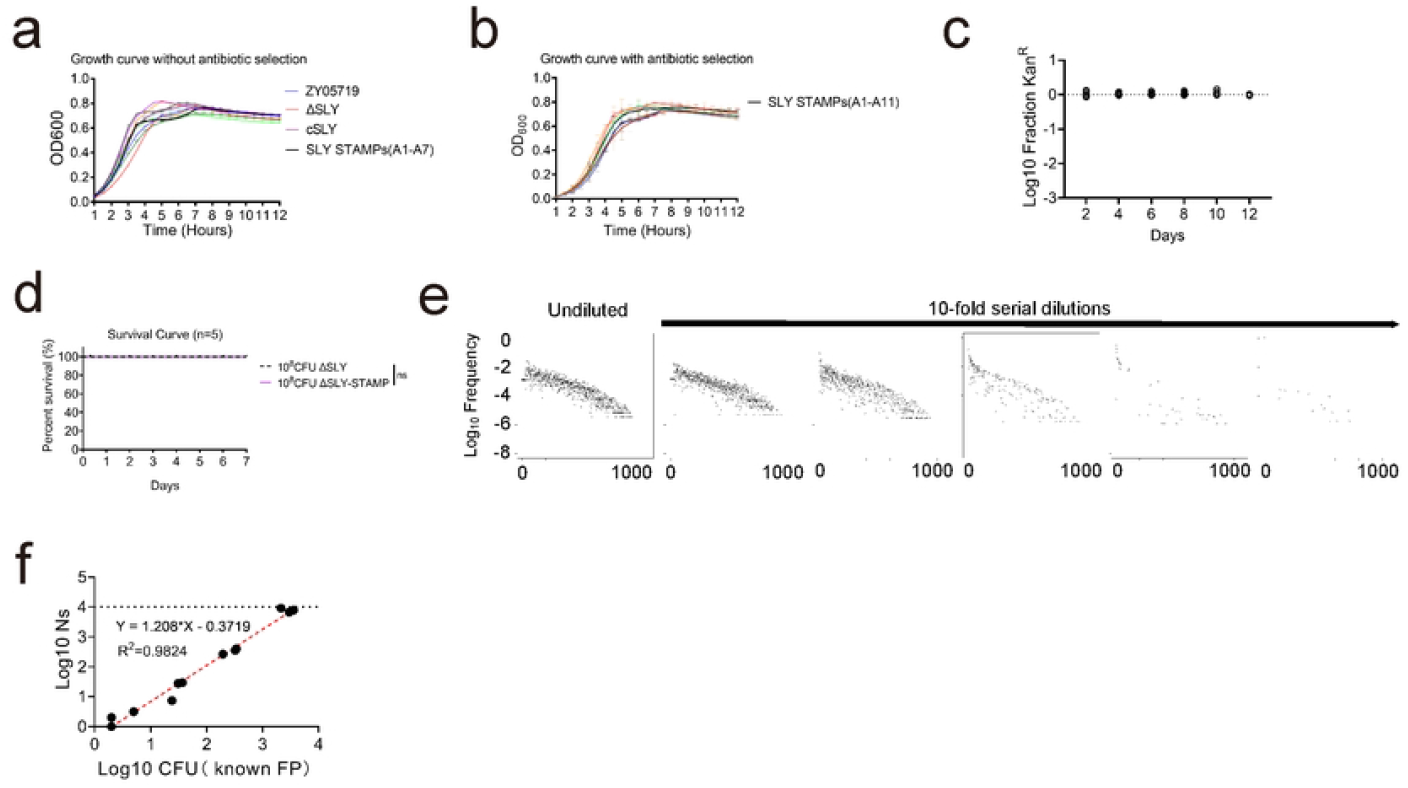
ΔSLY STAMP Library construction. (a) Growth curves of 7 ΔSLY barcoded clones, the wild type (ZY05719), ΔSLY and cSLY in THB media without Kan. (b) Growth curves of 11 ΔSLY barcoded clones in THB medium with kanamycin. No growth defects were observed with or without kanamycin. (c) The ΔSLY barcoded library was serially passaged in THB medium without Kan. Every two days, the fraction of cells containing barcodes were quantified as the fraction of kanamycin-resistant CFUs. All colonies retained the antibiotic resistance marker linked to barcode, indicating that the barcode is stable for at least 12 days without Kan. (d) Survival curves of mice challenged with 1×10^8^ CFU of ΔSLY or ΔSLY_-STAMP_. Mice received 1×10^8^ CFU were monitored for 7 days and both groups exhibited a 100% survival rate. (e, f) Sequencing the ΔSLY barcodes at various known bottleneck sizes. Ns accurately reflects founding FP values up to 10^4^. Survival curves were analyzed using the Log-rank (Mantel-Cox) test for (d): ns indicates no significant difference.

**Figure S5:**
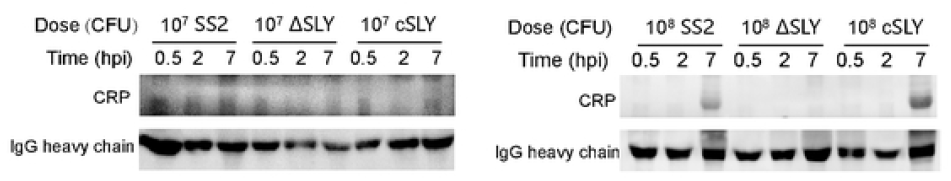
Western-blot detection of CRP in the serum of mice after SS2 or ΔSLY challenge. Immunoblot detection of CRP (25 kDa) and IgG heavy chain (50 kDa) in mouse serum. Mice were challenged with 1×10^8^ CFU of SS2, 1×10^8^ CFU of ΔSLY, and 1×10^8^ CFU of cSLY (complementation of ΔSLY) and euthanized at various time points (0.5 hpi, 2 hpi and 7 hpi). CRP bands were visible in the serum of mice challenged with 1×10^8^ CFU of SS2 WT and cSLY at 7 hpi.

## Notes

### Competing Interest Statement

The authors have declared no competing interest.

